# EphB2 and ERK signaling are required for heterotypic contact inhibition of locomotion to drive cell sorting

**DOI:** 10.1101/373696

**Authors:** Simon Brayford, Eduardo Serna-Morales, Andrei Luchici, Toru Hiratsuka, Brian M. Stramer

## Abstract

Interactions between different cell-types can induce distinct contact inhibition of locomotion (CIL) responses that are hypothesized to control population-wide behaviors during embryogenesis [1, 2]. However, our understanding of the signals that lead to cell-type specific repulsion, and the precise capacity of heterotypic CIL responses to drive emergent behaviors is lacking. Using a new *in vitro* model of heterotypic CIL between epithelial and mesenchymal cells, we show that fibrosarcoma cells (HT1080), but not fibroblasts (NIH3T3), are actively repelled by epithelial cells in culture. We show that knocking down EphB2 in fibrosarcoma cells specifically leads to disruption of the repulsion phase of CIL in response to interactions with epithelial cells. Furthermore, this heterotypic interaction requires ERK activation, downstream of EphB2 signaling. We also examine the population-wide effects when these various cell combinations, and their specific heterotypic CIL responses, are allowed to interact in culture. Mixtures of fibrosarcoma and epithelial cells – unlike fibroblasts and epithelial cells – lead to complete sorting and segregation of the two populations, and inhibiting their distinct CIL response by knocking down EphB2 or ERK in fibrosarcoma cells disrupts this emergent sorting behavior. Our understanding of the mechanisms underlying developmental behaviors such as cell sorting is lacking as predominant sorting hypotheses, such as differential adhesion, have recently been found inadequate in predicting the sorting of mesenchymal cells [3, 4]. These data suggest that heterotypic CIL responses, in conjunction with processes such as differential adhesion, may aid the sorting of cell populations during embryogenesis.

## Results and Discussion

### Fibroblasts and fibrosarcoma cells exhibit distinct responses upon collision with an epithelial cell monolayer

To study heterotypic cell-cell collisions, we developed a confrontation assay whereby two different cell-types are separated by a barrier, which upon removal, creates a uniform gap into which the different cell populations migrate and collide. Following a screen of a range of different epithelial versus mesenchymal cell-types, an interesting and unexpected phenomenon was revealed. When a population of migrating epithelial cells (HaCaT) encountered a population of migrating fibroblasts (NIH3T3), both populations ceased their forward migration, forming a sharp border (Figure 1A, B and Video S1). This is in stark contrast to fibrosarcoma cells (HT1080) which, upon collision with epithelial cells, seemed to undergo a complete repulsion (Figure 1A, B and Video S1). This result was not only seen with HaCaT epithelial cells, as fibrosarcoma collision with a human corneal epithelial cell line [5] also resulted in repulsion of the fibrosarcoma population (Figure S1). When no colliding partner was encountered, HaCaT epithelial cells migrated with a persistently increasing speed, which was similarly observed after their collision with fibrosarcoma cells, suggesting that epithelial cell motion was unaffected by their collision with the fibrosarcoma population (Figure 1C, D). In contrast, epithelial migration was severely reduced shortly after their collision with fibroblasts (Figure 1D). The repulsion of fibrosarcoma cells was further analyzed using particle image velocimetry (PIV) to track the entire cell population, which revealed a population-wide increase in the speed of the epithelial cells following collision with fibrosarcoma cells (Figure 1E and Video S2) showing that these cells have a distinct heterotypic response to epithelial cell collisions.

**Figure 1.**
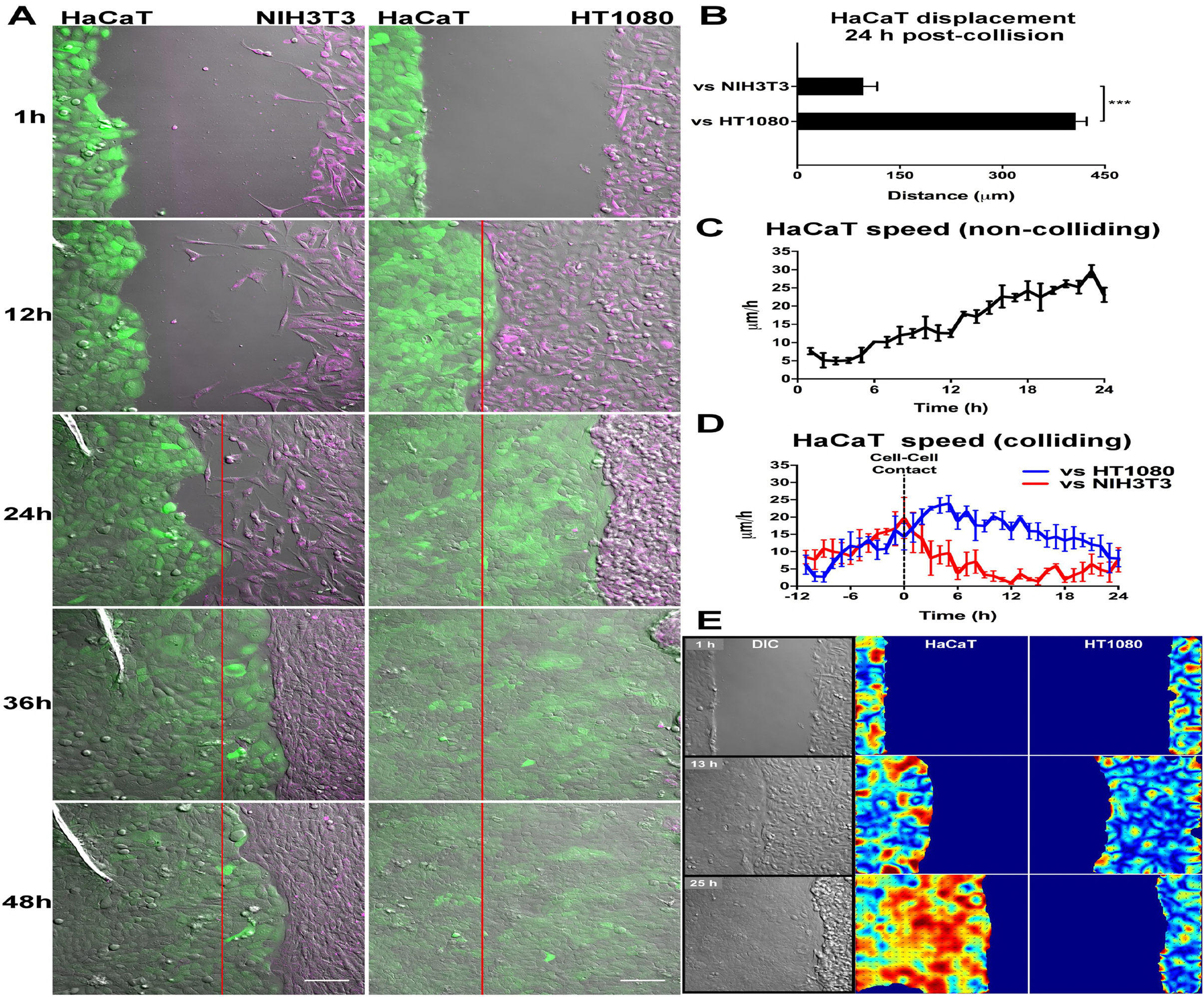
Fibroblasts and fibrosarcoma cells exhibit distinct responses upon collision with an epithelial cell monolayer. (A) Screenshots from representative movies of a confrontation assay in which epithelial cells (HaCaT) are allowed to collide with either fibroblasts (NIH3T3) or fibrosarcoma cells (HT1080). Epithelial cells (HaCaT) are labelled green and either fibroblasts (NIH3T3) or fibrosarcoma (HT1080) cells are labelled magenta. The red line indicates the position of initial cell-cell contact. Scale bars = 100 μm. (B) Quantification of HaCaT displacement 24 h after colliding with NIH3T3 or HT1080 cells in the confrontation assay (n = 3, error bars = SEM, ***P < 0.001, Student’s t-test). (C) Quantification of the speed of the front row of HaCaT cells throughout the confrontation assay when no other cell population is encountered, showing a persistent increase over 24 h (n = 3, error bars = SEM). (D) Quantification of the speed of the front row of HaCaT cells throughout the confrontation assay involving collision with either NIH3T3 or HT1080 cell populations. Note that the dotted line represents the time at which cell populations collide (n = 3, error bars = SEM). (E) Particle Image Velocimetry (PIV) analysis of the HaCaT and HT1080 interaction in ‘A’ showing an increase in speed of the entire population of HaCaT cells despite having contacted HT1080 cells. Blue to red represents a shift from low to high speed.

### Fibrosarcoma cells undergo active repulsion from epithelial cells

In order to determine whether the repulsion of fibrosarcoma cells by epithelial cells was an active process, we generated time-lapse movies of individual mesenchymal cells undergoing collisions with epithelial monolayers (Video S3) and examined their repulsive dynamics. An analysis of acceleration changes around collisions revealed that when fibrosarcoma cells collided with epithelial cells (i.e. heterotypic collisions), there was a sudden backward acceleration with respect to the colliding partner (Figure 2A) suggesting a sudden change in cell motion. This response was similar to previously reported homotypic collisions between *Drosophila* macrophages in vivo, which also undergo a classical CIL response involving active repulsion [6, 7]. The backward acceleration of fibrosarcoma cells was accompanied by a shift in the direction of their velocities before, during, and after the collision as the fibrosarcoma cells were repelled and migrated away from the epithelial cells (Figure 2B). In contrast, this repulsion was not observed when fibroblasts collided with epithelial cells, nor during fibrosarcoma-fibrosarcoma cell collisions (i.e. homotypic collisions), which led to cells continuing to migrate toward the colliding partner after collision (Figure 2A and B). Furthermore, quantification of nuclear distance from the point of collision over time revealed that heterotypic fibrosarcoma collisions led to a slowing of their motility before migrating away from epithelial cells, in contrast to homotypic collisions, which led to their continued forward motion (Figure 2C). These data highlight that fibrosarcoma cells show distinct CIL dynamics in response to collision with epithelial cells, which involves active cell repulsion.

**Figure 2.**
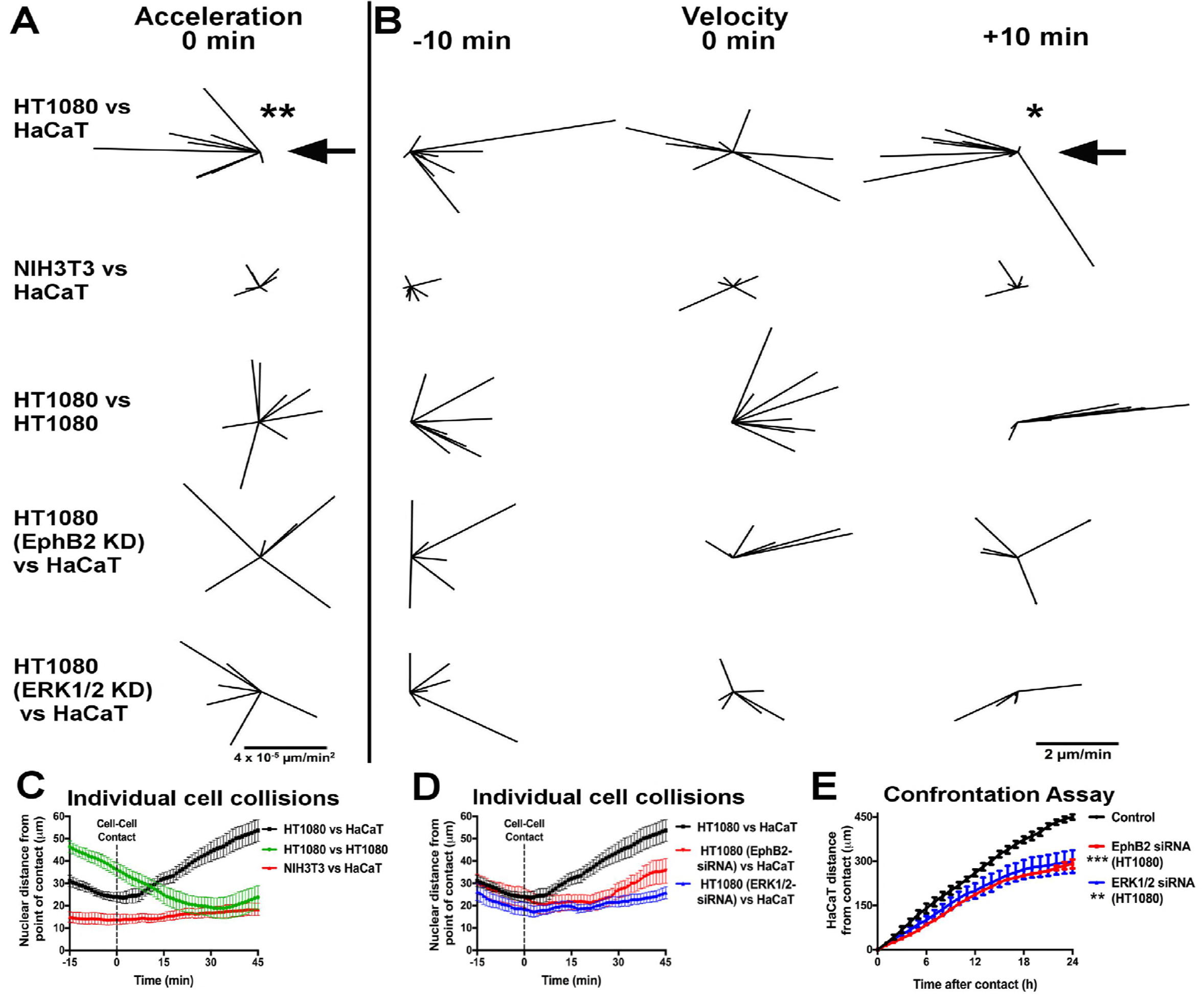
Fibrosarcoma cells undergo active repulsion upon collision with epithelial cells which is perturbed by EphB2 or ERK1/2 knockdown. (A) Vectors depicting the acceleration changes upon collision (time = 0 min) analysed using 10-minute time steps and normalized to the position of the colliding partner (denoted by the large arrow). A significant rearward acceleration is only observed in the fibrosarcoma (HT1080) vs epithelial (HaCaT) collision. (n ≥ 8 collisions per condition. **P < 0.01, non-parametric Wilcoxon signed rank test). (B) Vectors depicting the velocity of colliding cells at 10 mins before collision, at time of collision (0 min), and at 10 mins after collision, analysed using 5-minute time steps and normalized to the position of the colliding partner (denoted by the large arrow). A significant post-collision rearward velocity away from the colliding partner is only observed in the HT1080 vs HaCaT response. (n ≥ 8 collisions per condition. *P < 0.05, non-parametric Wilcoxon signed rank test). (C) Quantification of the distance of the nucleus from the position at which cell-cell contact occurs during heterotypic and homotypic interactions between HT1080, NIH3T3, and HaCaT cells. Note that after collision only heterotypic interaction between HT1080 and HaCaT cells leads to a rapid increase in cell separation (n ≥ 8 collisions per condition, error bars = SEM). (D) Quantification of the distance of the nucleus from the position at which cell-cell contact occurs during heterotypic HaCaT/HT1080 collisions in EphB2 or ERK1/2 knockdown in HT1080 cells. Note that a reduction in EphB2 or ERK1/2 reduces cell separation following collision (n ≥ 6 collisions per condition, error bars = SEM). (E) Quantification of the displacement of the front row of HaCaT cells after collision with HT1080 cells in confrontation assays comparing control, and EphB2 or ERK1/2 HT1080 knockdown. Note that the reduction in EphB2 or ERK1/2 in HT1080 cells slows the migration of the HaCaT population. (n = 3, error bars = SEM, ***P < 0.001, **P < 0.01, Friedman test).

### ERK activation downstream of EphB2 induces heterotypic repulsion of fibrosarcoma cells

There is much evidence in the literature for Eph receptors playing a role in epithelial cell interactions. EphA2 for example has been shown to drive the segregation of Ras-transformed epithelial cells from normal neighbors [8] and HEK293 cells overexpressing EphB2 have been shown to segregate from the same cells overexpressing ephrin-B1 [9]. However, less is known about whether ephrin signaling may control the repulsion and segregation of mesenchymal cell populations, similar to epithelial monolayers. Indeed, HT1080 fibrosarcoma cells express the Eph receptor EphB2, knockdown of which in these cells (Figure S2A) abolished the backward acceleration upon their collision with epithelial cells (Figure 2A). Furthermore, knockdown of EphB2 in fibrosarcoma cells led to a random distribution of the direction of velocities after collision suggesting that cells were randomly migrating away from epithelial cells rather than being actively repelled. (Figure 2B). Similarly, plotting nuclear distance from the point of collision over time revealed that EphB2 knockdown in fibrosarcoma cells slowed their separation from epithelial cells further showing that the repulsion phase of CIL was disrupted (Figure 2D). We also tested the effect of knocking down EphB2 in the confrontation assay and found that indeed, collective repulsion of fibrosarcoma cells following collision with a monolayer of epithelial cells was also disrupted (Figures 2E and S2B).

One signaling pathway previously reported to be controlled by Eph receptor activation is the ERK pathway. However, there is conflicting data for the role of ERK signaling specifically downstream of Eph receptor activation, with some reports suggesting an inhibition of ERK activity [10, 11] and others reporting an increase in ERK activity following Eph receptor stimulation [12-14]. We found that knockdown of ERK1/2 in fibrosarcoma cells (Figure S2C) phenocopied the results of EphB2 knockdown both in single-cell collisions (Figure 2A, B and D) and in confrontation assays (Figures 2E and S2B), showing that a combination of EphB2 activation and ERK signaling is required for fibrosarcoma repulsion from epithelial cells. In order to confirm that the effects of knocking down EphB2 or ERK signaling were indeed related to cell collisions and not general migration defects, we measured the speed and persistence of freely moving, non-colliding fibrosarcoma cells and found no significant difference in either speed (Figure S2D) or persistence (Figure S2E) in EphB2 or ERK knockdown cells suggesting that knockdown only affected events related to contact with another cell, rather than general migration. Interestingly, myosin II, which has been reported to be downstream of ERK signaling [15-17] and involved in Eph-mediated repulsion of other cell-types [18] did not appear to play a role in fibrosarcoma repulsion from epithelial cells as myosin inhibition with Blebbistatin had no effect in the confrontation assay (Figure S3).

We next examined ERK activity during heterotypic collisions using a mixing assay whereby two cell-types were combined 1:1, seeded together, and allowed to interact over time. Lysates from mixed-cell populations were then collected after 24 h and examined for ERK activity. Western blot analysis of these lysates revealed that phospho-ERK (pERK) was highest among epithelial-fibrosarcoma co-cultures, which was increased compared with either of these cell types individually (Figure 3A) suggesting that one population was contributing to ERK activation in the other. We also found that EphB2 knockdown in fibrosarcoma cells, whilst not affecting total ERK levels (Figure 3B), resulted in a significant decrease in levels of pERK in epithelial-fibrosarcoma co-cultures (Figure 3C), suggesting that ERK activation is downstream of EphB2 in the fibrosarcoma population.

**Figure 3.**
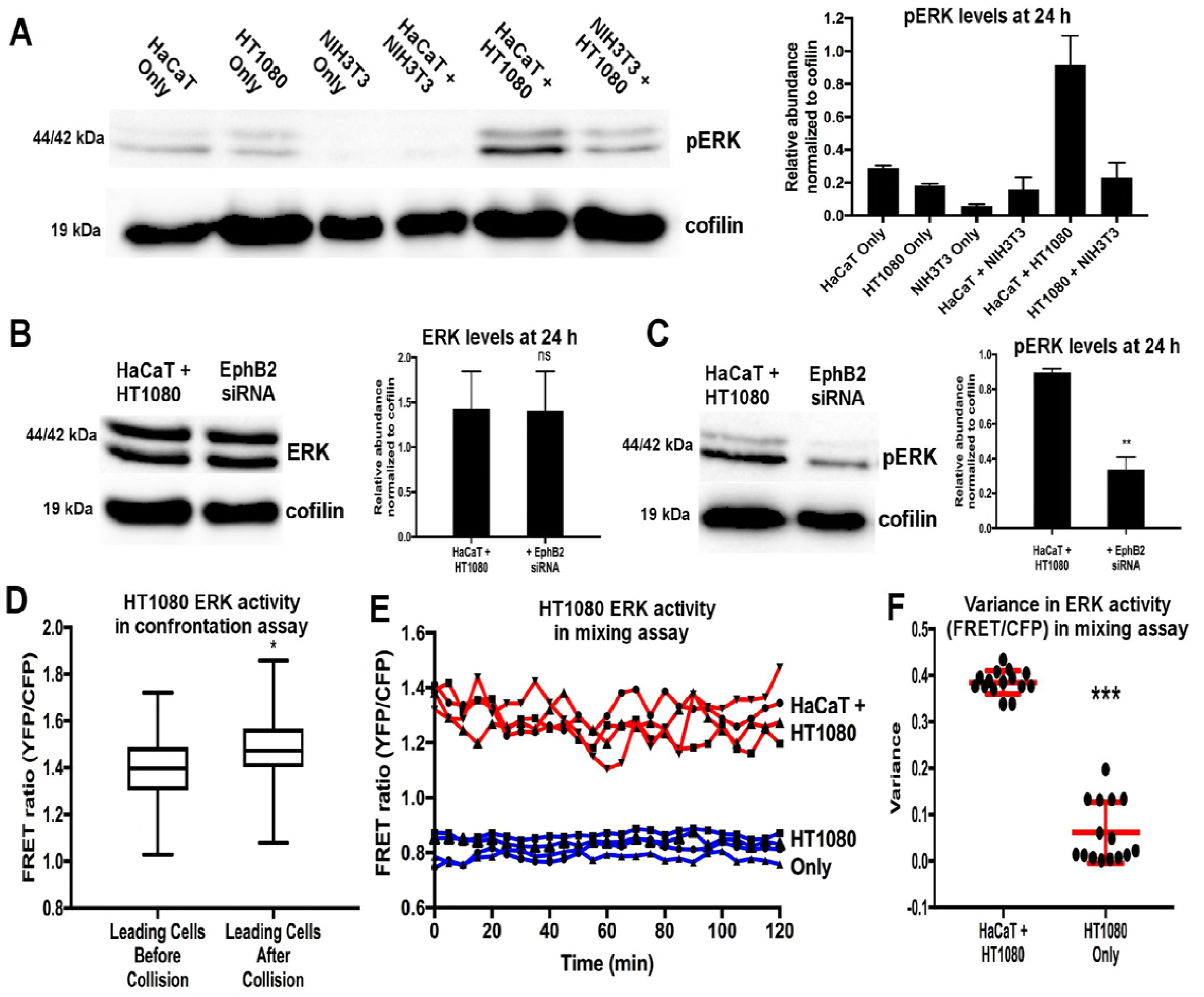
ERK activation is elevated in collisions between epithelial and fibrosarcoma cells. (A) Western blot for pERK in cell lysates collected at 24 h after plating cells individually or mixing the different cell types at a 1:1 concentration. The right panel shows a quantification of pERK levels revealing ERK activity is highest during co-culture of epithelial (HaCaT) and fibrosarcoma (HT1080) cells. (n = 3, error bars = SEM). (B) Western blot for total ERK in cell lysates in HaCaT/HT1080 co-cultures comparing controls and EphB2 knockdown in the HT1080 population. The right panel shows a quantification of total ERK levels revealing total ERK is unaffected by EphB2 knockdown in HT1080 cells. (n = 3, error bars = SEM, ns = not statistically significant, Student’s t-test). (C) Western blot for pERK in cell lysates in HaCaT/HT1080 co-cultures comparing controls and EphB2 knockdown in the HT1080 population. The right panel shows a quantification of pERK revealing ERK activity is reduced in the co-culture after EphB2 knockdown in HT1080 cells. (n = 3, error bars = SEM, **P < 0.01, Student’s t-test). (D) Quantification of ERK activity using the ERK FRET biosensor in the leading row of HT1080 cells in the confrontation assay 30 mins before collision vs 30 mins after collision. (n = 28 cells from 4 separate movies, boxplots represent range, median and quartiles, *P < 0.05, Mann-Whitney test). (E) Quantification of ERK activity from four representative tracks of HT1080 cells cultured alone (blue) or during co-culture with HaCaT cells (red). Note the increase in FRET activity in the co-cultured HT1080 population along with the increased fluctuation. (F) Quantification of the variance in ERK activity in randomly selected tracks of HT1080 cells cultured alone or during co-culture with HaCaT cells. (n = 15 tracks, line = mean, error bars ± SD, ***P < 0.001, Mann-Whitney test).

While western blotting shows pERK levels in an entire cell population, it provides no information as to ERK activation dynamics during individual cell-cell interactions. For this we used a FRET biosensor which has previously been demonstrated to report ERK activity in living cells [19]. In addition, this technique allows measurement of ERK activity in specific subpopulations of cells where western blotting does not. Following delivery of the biosensor into the fibrosarcoma population we repeated the confrontation assay and found that in fibrosarcoma cells at the migrating front, ERK activity was significantly increased after collision with epithelial cells (Figure 3D). We next analyzed time-lapse movies of the mixing assay and found an overall higher level of active ERK among fibrosarcoma cells in the presence of epithelial cells compared to fibrosarcoma cells alone (Figure 3E). Finally, we found that cells in the mixed group had a higher fluctuation in their ERK activity over time (by measuring variance in ERK activity of cell tracks) compared to fibrosarcoma only (Figure 3F). This was likely due to fibrosarcoma cells constantly undergoing heterotypic collisions with epithelial cells throughout the course of the assay. Taken together, these data suggest that ERK signaling acts downstream of EphB2 and is involved in the repulsion of fibrosarcoma cells upon their contact with epithelial cells.

### Fibrosarcoma and epithelial cell populations sort in culture through CIL interactions

To examine how distinct heterotypic and homotypic CIL dynamics affects the behavior of a mixed population of cells, we repeated the mixing assay and examined the population dynamics by generating time-lapse movies. We found that epithelial cells and fibroblasts mixed to form a homogenous, interspersed population, in contrast to epithelial and fibrosarcoma cells, which immediately began to sort from one another (Figure 4A and Video S4). In addition, automated tracking of the mesenchymal cells in these movies revealed a streaming-type behavior specifically among fibrosarcoma cells (Figure 4B and Video S4), highlighting this population’s segregation from epithelial cells over time. To examine whether the sorting behavior of fibrosarcoma cells was due in part to their collective migration, we examined the correlation in the direction of local instantaneous cell velocities, which measures the coupling of motion among neighboring cells in the population. This revealed that, while the local cell velocities of fibrosarcoma cells were more highly correlated than fibroblasts, fibrosarcoma cells still did not display a strong correlation in their motion (fibrosarcoma velocity correlation = 0.16 ± 0.01; fibroblast velocity correlation = 0.08 ± 0.01; mean ± SEM; 1 = perfect correlation, −1 = perfect anti-correlation) (Figure S4A and B). This suggests that, rather than physically following one another, fibrosarcoma cells are likely segregating from epithelial cells through some other mechanism.

**Figure 4.**
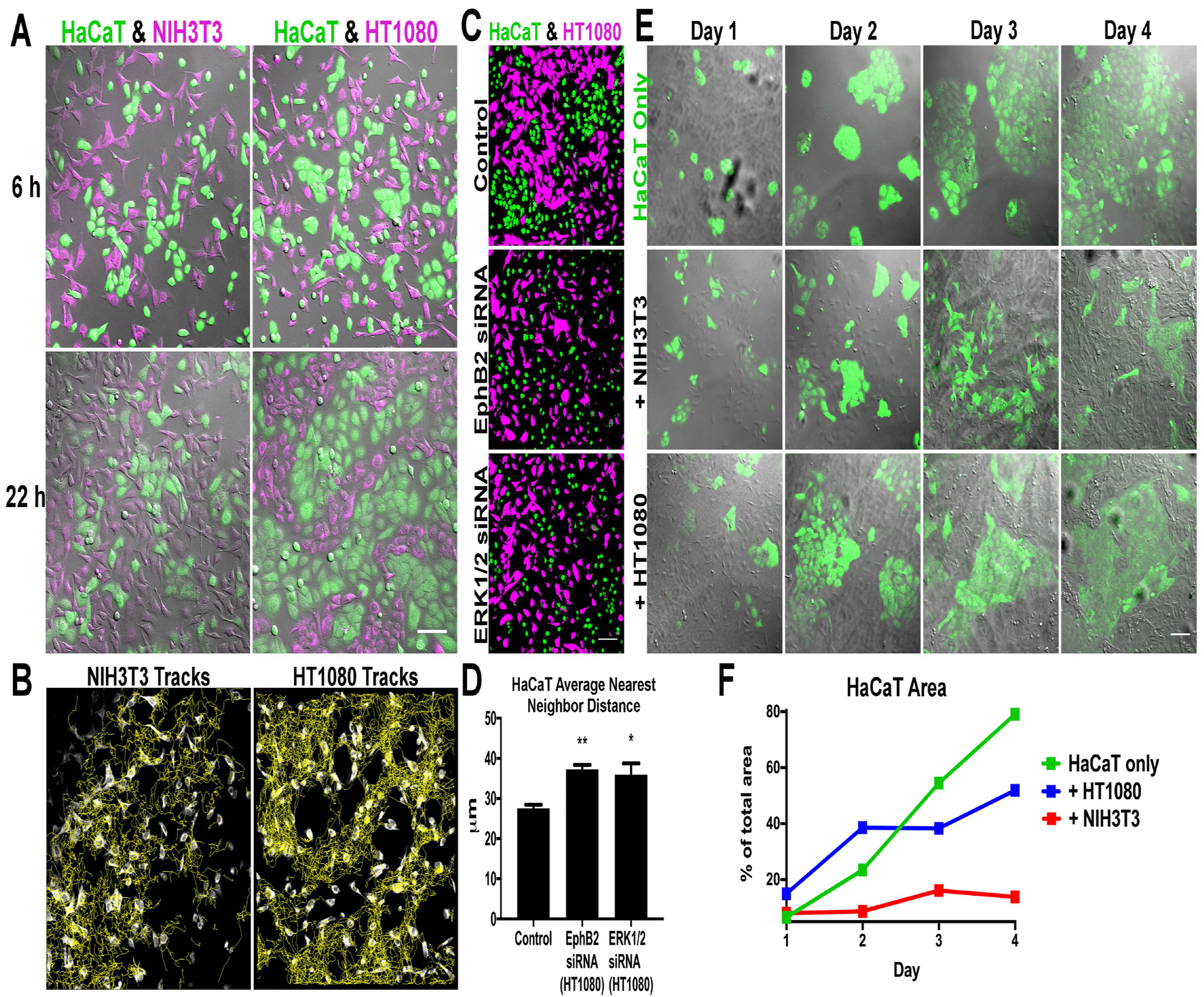
Fibrosarcoma cells segregate from epithelial cells in culture, a behavior which is disrupted by EphB2 or ERK knockdown. (A) Screenshots from movies of co-cultures of epithelial cells (HaCaT) with either fibroblasts (NIH3T3) or fibrosarcoma cells (HT1080). HaCaT cells are represented in green and mesenchymal cells in magenta. Scale bar = 100 μm. (B) Tracks of NIH3T3 or HT1080 cells throughout the movies in ‘A’ and Video S4. (C) Images of co-cultures (fixed at 24 h) of HaCaT cells (green) with either control, EphB2 or ERK1/2 knockdown HT1080 cells (magenta). Scale Bar = 100 μm. (D) The dispersion of HaCaT cells in ‘E’ quantified by measuring their distribution of nearest neighbor distances. An increase in HaCaT dispersion represents a reduction in their segregation from HT1080 cells. (n = 3, error bars = SEM, **P < 0.01, *P < 0.05, Student’s t-test). (E) Long term culture of HaCaT cells (green) alone or during co-culture with NIH3T3 or HT1080 cells (unlabeled). Note the specific increase in cluster size of HaCaT cells cultured alone or in the presence of HT1080 cells. Scale bar = 100 μm. (F) Quantification of area occupied by HaCaT cells from the images in ‘C’. Note that HaCaT cells are prevented from expanding specifically when co-cultured with NIH3T3 cells.

In order to examine whether the specific repulsive CIL dynamics of fibrosarcoma cells in response to epithelial cells was driving their segregation we inhibited Eph and ERK signaling in the fibrosarcoma population. Indeed, knockdown of EphB2 or ERK1/2 in fibrosarcoma cells resulted in a disruption to their segregation from epithelial cells as quantified by measuring the overall dispersion of epithelial cells (Figure 4C and D). Similarly, treatment with the MEK inhibitor, U0126, also resulted in disruption to the segregation between fibrosarcoma and epithelial cells (Figure S4C and E) further supporting a role for ERK signaling in the repulsion and segregation of these two populations. These data suggest that segregation of fibrosarcoma from epithelial cells is driven by the repulsive interaction between these two cell populations.

Finally, we investigated the phenomenon of cell-sorting over a longer time-course to examine the final patterns of the cell populations. When epithelial cells were plated sparsely and grown over a four-day period, they grew in clusters that eventually merged to form large, sheet-like structures (Figure 4E). However, epithelial cells grown in the presence of fibroblasts resulted in much smaller clusters with irregular shapes, as the two cell populations reached equilibrium (Figure 4E). In contrast, when grown together with fibrosarcoma cells, epithelial cells formed large colonies, seemingly unimpeded by the presence of fibrosarcoma cells (Figure 4E). Indeed, quantification of epithelial cell area over time revealed that their proliferation was specifically reduced in the presence of fibroblasts suggesting that epithelial cells may undergo contact inhibition of proliferation in the absence of a repulsive heterotypic CIL response (Figure 4F). We found no appreciable difference in basal proliferation rate between fibroblasts and fibrosarcoma cells (Figure S4F), suggesting that differential CIL dynamics was allowing fibroblasts, but not fibrosarcoma cells to out-compete epithelial cells for space, rather than a difference in their respective proliferation rates. Importantly, the ability of fibrosarcoma cells to almost completely segregate from epithelial cells differs from other 2D models of epithelial sorting in which epithelial cells do not undergo complete segregation, but rather lead to an intermediate aspect of separation that locks in small clusters of cells [9, 20, 21].

The differential adhesion hypothesis (DAH) has been the predominant model to explain the sorting of cell populations during development. Two different cell types can theoretically segregate simply by taking into account their differential adhesion (and differential surface tension) [22]. This model assumes that the cells in question segregate in a liquid-like phase separation guided by the reduction in adhesive-free energy as the two cell-types maximize their cell-cell adhesions. The problem with this concept in isolation is that we now know that groups of cells do not behave like perfect liquids; clusters of cells can quickly go from a fluid-like to a solid-like state as cell density increases in a process termed ‘jamming’, which leads to cells rapidly slowing their motion [23]. Therefore, DAH can only work in a regime where the cells are not proliferating and at a sweet spot in their density that allows sufficient fluidity to the tissue. Otherwise the cells will not be allowed to test the adhesive landscape sufficiently enough to lead to complete segregation. This is likely to be particularly important for epithelial populations (the predominant cell-type experimentally tested during cell sorting), which are not highly motile due to their strong cell-cell adhesions.

Here we show that a mesenchymal cell-type (HT1080) can rapidly and almost completely segregate from an epithelial population in 2D culture. Differential adhesion in this case is still playing a role in the separation due to the high surface tension of the epithelial population and the nearly non-existent mutual adhesion of the fibrosarcoma cells leading to the less cohesive cell (HT1080) surrounding the more cohesive cell type (HaCaT) as expected by the DAH [22]. However, DAH is not sufficient on its own; inhibiting the repulsion between these two cell types prevents their sorting behavior. We hypothesize that the strong heterotypic CIL response between fibrosarcoma and epithelial cells essentially helps fluidize the population allowing the cells to sample their adhesive surroundings. Furthermore, there also appears to be an aspect of fibrosarcoma CIL relieving contact inhibition of proliferation of the epithelial cells (possibly through a reduction in spatial constraints on the epithelial clusters) allowing epithelial colonies to grow and merge, which would also enhance the fluidity of the population. It is interesting to note that the other mesenchymal cell-type (NIH3T3) failed to segregate from the epithelial population as one would expect for a mixture of highly cohesive and non-cohesive cells [22]. In the fibroblast-epithelial combination, the epithelial cells are highly inhibited in their proliferation (likely due to a lack of heterotypic repulsion), which would enhance the spatial constraint on the epithelial colonies. In this case the lack of a repulsive CIL response prevents the epithelial colonies from growing enough to merge with neighboring colonies and thus separate from the fibroblast population. It is therefore possible that we are also highlighting an unexplored connection between contact inhibition of locomotion and contact inhibition of proliferation in this model despite the fact that these two processes are largely thought to be distinct behaviors [1].

CIL in recent years has been shown to play a role in the migration and patterning of numerous cell types during development. These data suggest that a repulsive CIL response in conjunction with other behaviors, such as differential adhesion, may also enhance the sorting of cell populations during embryogenesis.

## Acknowledgements

We thank Vicky Sanz-Moreno, Matthias Krause and Stephen Terry for use of cell-lines as well as Fiona Watt and Michiyuki Matsuda for use of and assistance with the ERK FRET biosensor. We would also like to thank Claudia Linker for feedback in writing the manuscript. This project has received funding from the European Research Council (ERC) under the European Union’s Horizon 2020 research and innovation program grant agreement No. 681808. TH has received funding from the European Union’s Horizon 2020 research and innovation program under the Marie Sklodowska-Curie grant agreement No. 704587.

## Author Contributions

Conceptualization, S.B. and B.M.S.; Methodology, S.B.; Software, E.S-M. and A.L.; Formal Analysis, S.B., E.S-M. and T.H.; Investigation, S.B.; Writing – Original Draft, S.B.; Writing – Review & Editing, S.B. and B.M.S.; Project Administration, B.M.S.; Funding Acquisition, B.M.S.

## Materials & Methods

### Cell culture

Immortalized human keratinocytes (HaCaT), immortalized mouse fibroblasts (NIH3T3), human fibrosarcoma (HT1080) and immortalized human corneal epithelial cells [5] were all maintained in Dulbecco’s Modified Eagle’s Medium (DMEM) with 4500 mg / L glucose, supplemented with 10 % FBS, at 37° C and 5 % CO_2_. For routine maintenance, cells were cultured in T75 plastic cell culture flasks and split via trypsinization approximately every 3 days or when approaching confluence.

### Cell labelling

CellTracker Green CMFDA and CellTracker Red CMTPX dyes (Invitrogen, Carlsbad, CA) were used to differentially label cell-types in mixed-cell assays. Cells were exposed to either dye at 1 μM in serum-free medium for 30 mins at 37° C before being washed once with PBS, trypsinized, re-suspended in complete medium and counted for downstream purpose.

### Live imaging

Movies were acquired using an LSM 880 inverted confocal microscope (Zeiss, Oberkochen, Germany). Cells were maintained at 37° C and 5 % CO_2_ for the duration of live imaging. Images were acquired using differential interference contrast (DIC) imaging along with airyscan filter sets for 488 or 561 lasers with either a 40X (NA 0.95) oil or 20X (NA 0.45) air objective (Zeiss).

### Confrontation assay

2-well culture inserts (Ibidi, Germany) were placed into the centre of the well of a 24 well μ-Plate (Ibidi, Germany). Two different pre-labelled cell-types were seeded into opposite chambers at a density of 1 × 10^5^ cells / cm^2^ and incubated overnight. The culture insert was then carefully removed and the well topped up with 1 mL complete culture medium before live imaging of the resulting 500 μm gap.

### Particle image velocimetry (PIV)

Time-lapse images of HaCaT and HT1080 cells from the confrontation assay movie (presented in Figure 1A) were manually segmented prior to analysis and pseudo-speckle analysis was performed as described previously [24]. For pseudo-speckle analysis, the size of the search image was chosen such that it spanned the maximum expected displacement of the cells during the acquisition time. To cover the whole search image, a cross-correlation coefficient was computed between source image and a sub-image of the search image shifted by one pixel. To remove anomalous tracking data, only displacements that had a cross-correlation coefficient above a certain threshold were kept. Finally, a spatial convolution with a Gaussian kernel and temporal convolution were used to interpolate the measured displacements to cover all the pixels within the frame. The complete algorithm for this analysis, including the filtering and interpolation was implemented in MatLAB (MathWorks^®^).

### Individual cell collisions and manual tracking

Analysis of single-cell collisions was carried out by mixing pre-labelled cells at low density, plating sparsely and allowing to adhere 4 h before imaging every 1 min at 20X. For the kinematics analysis, a collision was defined as the moment the the cell in question contacts any part of the colliding partner. The centroid of the cell nucleus, which could be easily identified from DIC movies was manually tracked using the mTrackJ plugin for ImageJ (NIH) to calculate the x,y coordinates of the cell at all time-points.

### Kinematics analysis

Kinematics analysis of the velocity and acceleration of cells was calculated as previously described [6, 25]. In order to assess the statistical significance of the direction of cells after collision, a binomial test with a probability of success of 95% was performed on the cell velocity unit vectors every five minutes from 5 min before to 20 min after collision. To assess the statistical significance of acceleration, a one-sample t test of the horizontal component of the vectors was performed. This test analyzes how far from zero the displacement average of the horizontal component of the vectors is at the collision time. If the average is similar to zero it suggests that the displacements are random.

### Gene silencing by small interfering RNA (siRNA)

HT1080 cells were plated onto 6-well plates at 2 × 10^5^ cells / well and allowed to attach overnight. Cells were transfected with pre-validated siRNA sequences to knockdown human EphB2 (Sigma, cat. no. EHU060511) or human p44/42 MAPK (ERK1/2) (Cell Signaling Technologies, Cat no. 6560). Mock transfection was used as a negative control. siRNA was transfected using Oligofectamine^®^ reagent (Invitrogen) according to the manufacturer’s instructions. Experiments were carried out 48 h post-transfection.

### Western blotting and antibodies

Total cellular proteins from individual cells or mixed-cell populations were prepared by rinsing cells with cold PBS and scraping with RIPA buffer (20 mM Tris pH 7.4, 150 mM sodium chloride, 1% (v/v) Nonidet P-40, 0.5% (w/v) sodium deoxycholate, 1 mM EDTA, 0.1% (w/v) SDS) in the presence of protease and phosphatase inhibitor cocktails (Roche Diagnostics, Indianapolis, IN). Approximately equal amounts of protein (20 μg) were resolved on 12.5% SDS-PAGE gels before electro-transfer to PVDF membranes. Following blocking in 5% (w/v) BSA in TBST, immunoblotting was performed using anti-EphB2 (CST, #83029), anti-GADPH (Millipore, #ABS16), anti-p44/42 MAPK (CST #9102), anti-cofilin (CST #5175) or anti-Phospho-p44/42 MAPK to detect EphB2, GAPDH, ERK1/2, cofilin and pERK1/2 (Thr202/Tyr204) protein respectively. The membranes were then washed with TBST and incubated with species appropriate HRP-conjugated IgG secondary antibodies (Agilent Technologies, Santa Clara, CA). Chemiluminescence was measured using ImageJ (NIH) after applying Clarity^™^ Western ECL substrate (BioRad, Hercules, CA).

### ERK FRET biosensor plasmid

The lentiviral plasmid of nucleus-localized FRET biosensor for ERK (EKAREV-NLS) has been previously characterized [19] and was a gift from Dr. Michiyuki Matsuda at Kyoto University, Japan.

### Lentiviral transduction

EKAREV-NLS ERK biosensor was expressed in HT1080 cells by lentiviral transduction. EKAREV-ELS in replication-defective, self-inactivating lentiviral pCSII vector was co-transfected with packaging plasmid (pCAG-HIVgp) and VSV-G-/Rev-expressing plasmid (pCMV-VSVG-RSV-Rev) into Lenti-X 293T cells (Clontech). High-titre viral solution was prepared and used for transduction into HT1080 cells.

### Cell mixing assay

Cells were pre-labelled with either cell-tracker red or green, counted and mixed in 1:1 suspensions so that 5 ×10^4^ of each cell type was seeded per well in a 24-well imaging μ-Plate (total of 1 × 10^5^ cells / well). Cells were allowed 6 h to adhere before being imaged overnight for a total of 22 h (for the wild-type analysis). For the RNAi mixing assay, cells were allowed to mix for 24 h before being fixed with 4% paraformaldehyde. All nuclei were then labelled with DRAQ5 (Invitrogen) before coverslips were mounted. Still images were then acquired and the green channel was used to create a mask over the HT1080 cells’ nuclei, leaving only the HaCaT cells’ nuclei visible and then using the nearest neighbour distance calculation in ImageJ (NIH).

### Automated cell tracking and alignment analysis

For automated tracking, the image channel corresponding to NIH3T3 or HT1080 cells was thresholded in ImageJ so that individual cells could be detected as particles in the TrackMate plugin and tracks generated. The nearest neighbor analysis obtains the correlation between the instant displacement vectors of pairs of nearest neighbor cells at every frame. First, pairs of nearest neighbors are found, afterwards, each cell is tracked onto the next frame to generate an instant displacement vector. Posteriorly, a Pearson correlation is performed to find the linear correlation between the instant displacement vector of each pair of neighbors, which can be represented as the cosine of the angle formed by two vectors. To represent the correlation data, an average of the correlation coefficient of all pairs of nearest neighbors at each frame was performed.

### Long-term co-culture assay

HaCaT cells were pelleted by centrifugation and re-suspended in CellTrace^™^ CFSE (Invitrogen) at a working concentration of 5 μM in PBS and incubated for 20 minutes at 37°C. Cells were again pelleted and re-suspended in fresh culture medium to a density of 2 ×10^4^ cells / mL and incubated for 10 minutes to allow the CFSE reagent to undergo acetate hydrolysis. This suspension was then mixed with either DMEM (control), an NIH3T3 cell suspension (unlabelled) or HT1080 cell suspension (unlabelled) and added to a 24-well plate for a total of 20,000 cells per well and incubated for 4 days, fixing with 4% paraformaldehyde every 24 h and mounting coverslips with ProlongGold (Invitrogen). Coverslips were then imaged with a 20x objective for analysis of HaCaT growth area using ImageJ (NIH).

### Proliferation assay

NIH3T3 and HT1080 cells were seeded in a 96-well plate at 1000 cells per well and incubated for 4 h. Growth medium was then carefully removed and 1X CyQUANT^®^ NF (Invitrogen) dye reagent was added. Cells were then incubated at 37°C for 1 h. This incubation period is required for equilibration of dye–DNA binding, resulting in a stable fluorescence endpoint. The fluorescence intensity of each sample was then imaged every 24 h using a fluorescence microplate reader with excitation at ~485 nm and emission detection at ~530 nm. Fresh growth medium was added every 48 h. Cell numbers at each time-point were determined using a standard curve as per the manufacturer’s instructions.

### Drug Treatments

For the myosin II inhibition experiment, the photo-stable derivative of blebbistatin, (S)-nitro-Blebbistatin (Cayman Chemical, Ann Arbor, MI) was used at a final concentration of 50 μM in complete medium. For MEK inhibition experiments, U0126 (Merck, Kenilworth, NJ) was used at a final concentration of 20 μM in complete medium.

### Statistical analyses

Statistical analyses are described in each figure legend.

---

**Figure S1. Related to Figure 1**

(A) Screenshots from representative movies of a confrontation assay in which Human Corneal Epithelial Cells (HCE) are allowed to collide with fibrosarcoma cells (HT1080). HCE cells are labelled green and HT1080 cells are labelled magenta. Scale bar = 100 μm.

**Figure S2. Related to Figure 2**

(A) Western blot confirming successful knockdown of EphB2 protein in HT1080 cells 48 h post-treatment with siRNA compared with non-transfected cells (wild-type).

(B) Screenshots from representative movies of a confrontation assay in which epithelial cells (HaCaT) are allowed to collide with either control, EphB2 or ERK1/2 knockdown fibrosarcoma cells (HT1080). Epithelial cells (HaCaT) are labelled green and fibrosarcoma (HT1080) cells are labelled magenta. The red line indicates the position of initial cell-cell contact. Scale bars = 100 μm.

(C) Western blot confirming successful knockdown of ERK1/2 protein in HT1080 cells 48 h post-treatment with siRNA compared with non-transfected cells (wild-type).

(D) Quantification of non-colliding HT1080 cell speed in cells transfected with either EphB2 or ERK1/2 siRNA compared with non-transfected cells (wild-type). (n = 5-10 cells, error bars = SEM, ns = not statistically significant, Student’s t-test).

(E) Quantification of non-colliding HT1080 cell persistence in cells transfected with either EphB2 or ERK1/2 siRNA compared with non-transfected cells (wild-type). (n = 5-10 cells, error bars = SEM, ns = not statistically significant, Student’s t-test).

**Figure S3. Related to Figure 2**

(A) Screenshots from representative movies of a confrontation assay in which epithelial cells (HaCaT) are allowed to collide with fibrosarcoma cells (HT1080) after blebbistatin treatment to inhibit Myosin II. HaCaT cells are labelled green and HT1080 cells are labelled magenta. Scale bar = 100 μm.

(B) Quantification of the average speed of the HT1080 cell population in the confrontation assay before the collision occurs. Cells treated with Blebbistatin migrated with a higher average pre-collision speed than untreated cells, indicating that blebbistatin has been effective in myosin II inhibition (n = 3, error bars = SEM, P* < 0.05, Student’s t-test).

(C) Quantification of the displacement of the front row of HaCaT cells after collision with HT1080 cells in the confrontation assay comparing untreated control and blebbistatin treated cultures. (n = 3, error bars = SEM).

**Figure S4. Related to Figure 4**

(A) Directional vector-field plots for NIH3T3 and HT1080 cells (taken at time 15 h from the respective movies in Video S4).

(B) Quantification of the local alignment of cell velocities of NIH3T3 or HT1080 cells during co-culture with HaCaT cells. A positive correlation corresponds to polar alignment of velocities and a negative correlation an anti-parallel alignment. Note the higher correlation of velocity vectors in HT1080-HaCaT co-culture which is suggestive of streaming behavior. (n = 140 cells, line = mean, error bars ± SEM ***P < 0.001, Mann-Whitney test).

(C) Images of co-cultures of epithelial cells (HaCaT) with fibrosarcoma cells (HT1080) comparing DMSO control with U0126 treatment to inhibit ERK1/2 signaling.

(D) Quantification of HaCaT cell numbers at 6 and 24 h which indicates that U0126 inhibition of ERK does not selectively impact the growth rate of HaCaT cells. (n = 3, error bars = SEM, ns = not statistically significant, Student’s t-test).

(E) The dispersion of HaCaT cells quantified by measuring their distribution of nearest neighbor distances. An increase in HaCaT dispersion represents a reduction in their segregation from HT1080 cells. (n = 3, error bars = SEM, *P < 0.05, Student’s t-test).

(F) Basal proliferation rates of NIH3T3 and HT1080 cells measured over a 5-day period which demonstrates that rates do not differ between the two cell-types. (n = 3, error bars = mean ± SD).

**Video S1. Related to Figure 1**

Example movies of the confrontation assay following barrier removal. HaCaT epithelial cells are labelled green and either NIH3T3 fibroblasts or HT1080 fibrosarcoma cells are labelled magenta. Time steps = 1 h. Scale bar = 100 μm.

**Video S2. Related to Figure 1**

Particle Image Velocimetry (PIV) heat-map of the HaCaT and HT1080 interaction in Video S1 showing an increase in speed of the entire population of HaCaT cells after colliding with HT1080 cells. Blue to red represents a shift from low to high instantaneous velocity.

**Video S3. Related to Figure 2**

Examples of DIC movies showing individual HT1080 fibrosarcoma cells (Control, EphB2 or ERK1/2 knockdown) colliding with HaCaT epithelial cells or homotypic collisions between HT1080 cells. Red lines represent manual tracking used for the kinematics analysis presented in Figure 2. Time steps = 1 min. Scale Bar = 20 μm.

**Video S4. Related to Figure 4**

(Top) Example movies of the mixing assay. HaCaT epithelial cells are labelled in green, other cells magenta. Time steps = 10 min. Scale bar = 100 μm. (Bottom) Tracks of NIH3T3 fibroblasts (left) and HT1080 fibrosarcoma cells (right) from the movies above. Blue through red coloration of tracks represents temporal position in movie to show that, in the HT1080 case (bottom right), cells eventually become segregated from HaCaT cells (not labelled).

## References

1. Stramer, B., and Mayor, R. (2016). Mechanisms and in vivo functions of contact inhibition of locomotion. Nature reviews. Molecular cell biology.

2. Theveneau, E., Steventon, B., Scarpa, E., Garcia, S., Trepat, X., Streit, A., and Mayor, R. (2013). Chase-and-run between adjacent cell populations promotes directional collective migration. Nat Cell Biol 15, 763–772.

3. Pawlizak, S., Fritsch, A.W., Grosser, S., Ahrens, D., Thalheim, T., Riedel, S., Kießling, T.R., Oswald, L., Zink, M., Manning, M.L., et al. (2015). Testing the differential adhesion hypothesis across the epithelial−mesenchymal transition, (Bristol, UK: Institute of Physics Pub).

4. Tambe, D.T., and Fredberg, J.J. (2015). And I hope you like jamming too. New J Phys 17.

5. Terry, S.J., Dona, F., Osenberg, P., Carlton, J.G., and Eggert, U.S. (2018). Capping protein regulates actin dynamics during cytokinetic midbody maturation. Proc Natl Acad Sci U S A 115, 2138–2143.

6. Davis, J.R., Luchici, A., Mosis, F., Thackery, J., Salazar, J.A., Mao, Y., Dunn, G.A., Betz, T., Miodownik, M., and Stramer, B.M. (2015). Inter-cellular forces orchestrate contact inhibition of locomotion. Cell 161, 361–373.

7. Davis, J.R., Huang, C.Y., Zanet, J., Harrison, S., Rosten, E., Cox, S., Soong, D.Y., Dunn, G.A., and Stramer, B.M. (2012). Emergence of embryonic pattern through contact inhibition of locomotion. Development 139, 4555–4560.

8. Porazinski, S., de Navascues, J., Yako, Y., Hill, W., Jones, M.R., Maddison, R., Fujita, Y., and Hogan, C. (2016). EphA2 Drives the Segregation of Ras-Transformed Epithelial Cells from Normal Neighbors. Curr Biol 26, 3220–3229.

9. Poliakov, A., Cotrina, M.L., Pasini, A., and Wilkinson, D.G. (2008). Regulation of EphB2 activation and cell repulsion by feedback control of the MAPK pathway. J Cell Biol 183, 933–947.

10. Miao, H., Nickel, C.H., Cantley, L.G., Bruggeman, L.A., Bennardo, L.N., and Wang, B. (2003). EphA kinase activation regulates HGF-induced epithelial branching morphogenesis. J Cell Biol 162, 1281–1292.

11. Elowe, S., Holland, S.J., Kulkarni, S., and Pawson, T. (2001). Downregulation of the Ras-mitogen-activated protein kinase pathway by the EphB2 receptor tyrosine kinase is required for ephrin-induced neurite retraction. Mol Cell Biol 21, 7429–7441.

12. Pratt, R.L., and Kinch, M.S. (2002). Activation of the EphA2 tyrosine kinase stimulates the MAP/ERK kinase signaling cascade. Oncogene 21, 7690–7699.

13. Vindis, C., Cerretti, D.P., Daniel, T.O., and Huynh-Do, U. (2003). EphB1 recruits c-Src and p52Shc to activate MAPK/ERK and promote chemotaxis. J Cell Biol 162, 661–671.

14. Kandouz, M., Haidara, K., Zhao, J., Brisson, M.L., and Batist, G. (2010). The EphB2 tumor suppressor induces autophagic cell death via concomitant activation of the ERK1/2 and PI3K pathways. Cell Cycle 9, 398–407.

15. Aoki, K., Kondo, Y., Naoki, H., Hiratsuka, T., Itoh, R.E., and Matsuda, M. (2017). Propagating Wave of ERK Activation Orients Collective Cell Migration. Dev Cell 43, 305–317 e305.

16. Kamioka, Y., Sumiyama, K., Mizuno, R., Sakai, Y., Hirata, E., Kiyokawa, E., and Matsuda, M. (2012). Live imaging of protein kinase activities in transgenic mice expressing FRET biosensors. Cell Struct Funct 37, 65–73.

17. Mizuno, R., Kamioka, Y., Kabashima, K., Imajo, M., Sumiyama, K., Nakasho, E., Ito, T., Hamazaki, Y., Okuchi, Y., Sakai, Y., et al. (2014). In vivo imaging reveals PKA regulation of ERK activity during neutrophil recruitment to inflamed intestines. J Exp Med 211, 1123–1136.

18. Fagotto, F., Rohani, N., Touret, A.S., and Li, R. (2013). A molecular base for cell sorting at embryonic boundaries: contact inhibition of cadherin adhesion by ephrin/ Eph-dependent contractility. Dev Cell 27, 72–87.

19. Komatsu, N., Aoki, K., Yamada, M., Yukinaga, H., Fujita, Y., Kamioka, Y., and Matsuda, M. (2011). Development of an optimized backbone of FRET biosensors for kinases and GTPases. Mol Biol Cell 22, 4647–4656.

20. Taylor, H.B., Khuong, A., Wu, Z., Xu, Q., Morley, R., Gregory, L., Poliakov, A., Taylor, W.R., and Wilkinson, D.G. (2017). Cell segregation and border sharpening by Eph receptor-ephrin-mediated heterotypic repulsion. J R Soc Interface 14.

21. Taylor, W.R., Morley, R., Krasavin, A., Gregory, L., Wilkinson, D.G., and Poliakov, A. (2012). A mechanical model of cell segregation driven by differential adhesion. PLoS One 7, e43226.

22. Foty, R.A., and Steinberg, M.S. (2013). Differential adhesion in model systems. Wiley Interdiscip Rev Dev Biol 2, 631–645.

23. Sadati, M., Taheri Qazvini, N., Krishnan, R., Park, C.Y., and Fredberg, J.J. (2013). Collective migration and cell jamming. Differentiation 86, 121–125.

24. Betz, T., Koch, D., Lim, D., and Kas, J.A. (2009). Stochastic actin polymerization and steady retrograde flow determine growth cone advancement. Biophys J 96, 5130–5138.

25. Dunn, G.A., and Paddock, S.W. (1982). Analysing the motile behaviour of cells: a general approach with special reference to pairs of cells in collision. Philos Trans R Soc Lond B Biol Sci 299, 147–157.

